# Antenna modification leads to enhanced nitrogenase activity in a high light tolerant cyanobacterium

**DOI:** 10.1101/2021.11.15.468767

**Authors:** Anindita Bandyopadhyay, Zi Ye, Zuzana Benedikty, Martin Trtilek, Himadri B. Pakrasi

## Abstract

Biological nitrogen fixation is an energy intensive process that contributes significantly towards supporting life on this planet. Among nitrogen-fixing organisms, cyanobacteria remain unrivaled in their ability to fuel the energetically expensive nitrogenase reaction with photosynthetically harnessed solar energy. In heterocystous cyanobacteria light-driven, photosystem I (PSI)-mediated ATP synthesis plays a key role in propelling the nitrogenase reaction. Efficient light transfer to the photosystems rely on phycobilisomes (PBS), the major antenna protein complexes. PBS undergo degradation as a natural response to nitrogen starvation. Upon nitrogen availability, these proteins are resynthesized back to normal levels in vegetative cells, but their occurrence and function in heterocysts remains inconclusive. *Anabaena* 33047 is a heterocystous cyanobacterium that thrives under high light, harbors higher amounts of PBS in its heterocysts and fixes nitrogen at higher rates compared to other heterocystous cyanobacteria. To assess the relationship between PBS in heterocysts and nitrogenase function, we engineered a strain that retains high amounts of the antenna proteins in its heterocysts. Intriguingly, under high light intensities the engineered strain exhibited unusually high rates of nitrogenase activity compared to the wild type. Spectroscopic analysis revealed altered PSI kinetics in the mutant, with increased cyclic electron flow around PSI, a route that contributes to ATP generation and nitrogenase activity in heterocysts. Retaining higher levels of PBS in heterocysts appears to be an effective strategy to enhance nitrogenase function in cyanobacteria that are equipped with the machinery to operate under high light intensities.

**Importance:** The function of phycobilisomes, the large antenna protein complexes in heterocysts has long been debated. This study provides direct evidence of the involvement of these proteins in supporting nitrogenase activity in *Anabaena* 33047, a heterocystous cyanobacterium that has affinity for very high light intensities. This strain was previously known to be recalcitrant to genetic manipulation and hence despite its many appealing traits, remained largely unexplored. We developed a genetic modification system for this strain and generated a Δ*nblA* mutant that exhibited resistance to phycobilisome degradation upon nitrogen starvation. Physiological characterization of the strain indicated that PBS degradation is not essential for acclimation to nitrogen deficiency and retention of PBS is advantageous for nitrogenase function.

## Introduction

Nitrogen fixation is a crucial process by which molecular nitrogen is rendered accessible to all life forms. The process is energy intensive, making fixed nitrogen a sparse commodity in many natural habitats and cultivated lands. Of the nitrogen fixing organisms, photosynthetic prokaryotes with the inherent ability to couple the energy demanding nitrogenase reaction with energy generating photosynthesis have a distinct advantage. Cyanobacteria constitute a group of oxygenic photosynthetic prokaryotes, some members of which have the dual ability to fix carbon and nitrogen. Nitrogenase, the enzyme catalyzing microbial nitrogen fixation is prone to destruction by oxygen and as such, diazotrophic cyanobacterial species have evolved to separate the incompatible processes of photosynthesis and nitrogen fixation temporally or spatially.

Some filamentous cyanobacteria segregate nitrogen fixation in specialized cells called heterocysts. Heterocysts lack functional photosystem II (PSII) and oxygenic photosynthesis but have a fully functional photosystem I (PSI) that is involved in generating ATP by cyclic photophosphorylation (1). Studies have indicated that the main source of ATP for nitrogen fixation in heterocysts is the light reactions in the thylakoids and that the required ATP can be provided entirely by light driven cyclic photophosphorylation (2–4). Compared to adjoining vegetative cells, heterocysts have a significantly higher abundance of PSI protein subunits (5, 6) as well as subunits of the Cyt-b_6_f complex involved in cyclic electron transport (3, 7). In addition, the number of photosynthetic units (ratio of chlorophyll a/P700) per heterocyst is 50% higher compared to vegetative cells (2)7, (8). All these evidences point to a distinct role for light and PSI in heterocyst function.

Light harvest in cyanobacteria is achieved by large protein complexes called phycobilisomes (PBS) that transfer energy to PSI and PSII reaction centers(9). PBS undergo degradation as an adaptive strategy to various environmental stress conditions like high light and nitrogen deficiency (10, 11). Although the mechanism of PBS degradation is yet to be fully elucidated, it has been established that NblA, a small protein consisting of 60 amino acids is a key player in the process (12). The effect of *nblA* deletion has been extensively investigated in non-diazotrophic model strains of cyanobacteria in which PBS degradation is an essential adaptive mechanism to survive high light and nitrogen stress (13–15). In contrast, PBS degradation is less likely to have a role in acclimation to these stress factors in high light tolerant, diazotrophic cyanobacteria and the effect of *nblA* deletion in these strains is yet to be investigated.

*Anabaena* sp. ATCC 33047 is a fast growing filamentous cyanobacteria that has garnered interest for its ability to thrive under very high light intensities and fix carbon and nitrogen at high rates (16). This strain was shown to harbor significant amounts of PBS in its heterocysts, implying expensive re-synthesis of these antenna protein complexes after the initial nitrogen acclimation phase. However, the function of these proteins in mature heterocysts and its relation to nitrogen fixation was not obvious (17). Earlier studies implicating a role for PBS in nitrogen fixation, specifically in energy transfer to PSI (7, 18, 19), led us to develop a genome engineering strategy for this previously recalcitrant strain and generate a Δ*nblA* mutant that would enable us to investigate the effect of heterocyst antenna modification on nitrogenase function. The Δ*nblA* mutant retained high amounts of PBS in its heterocysts and exhibited 2-3-fold higher nitrogenase activity compared to the WT. Spectroscopic analysis of the mutant indicated higher cyclic electron flow, possibly resulting from higher energy transfer to PSI, facilitated by excess PBS in heterocysts. This in turn leads to higher ATP generation and enhanced nitrogenase activity. Our study suggests that augmenting light capture by heterocysts of high light tolerant cyanobacteria can be an effective strategy to enhance nitrogen fixation.

## Results

### *Anabaena* 33047 thrives under very high light intensities

*Anabaena* 33047 exhibited rapid growth both in the presence and absence of added nitrogen sources and an increase in growth rate was observed with increasing light intensity (Fig. 1 A, B). When grown at 42°C in media supplemented with nitrogen sources and 1% CO_2_, the fastest growth rate (doubling time of 3.18±.16 h) was observed under 1500 - 2000 μmol photons m^−2^s^−^1 of light (Fig. 1A). At higher light intensities (up to 3000 μmol photons m^−2^s^−1^), although no increase in growth rate was observed, higher biomass accumulation could be achieved. When grown under light intensities of 3000 μmol photons m^−2^s^−1^ high amounts of exopolysaccharide (EPS) was secreted into the media which led to excessive clumping of filaments (data not shown). When grown under 2000 μmol photons m^−2^s^−^1 of light in nitrogen-deplete media, a doubling time of 3.8±0.43 h was observed (Fig. 1B). These growth rates are considerably higher compared to the commonly studied model filamentous strain *Anabaena* 7120, which under 400 μmol photons m^−2^s^−1^ light at 28°C exhibits a doubling time of 8.5±0.32 h under N_2_-sufficient and 12.2±0.86 h under N_2_-deficient conditions (Fig S2). Under light intensities >1500 μmol photons m^−2^s^−1^ light this strain exhibits growth inhibition (data not shown). In contrast, growth of *Anabaena* 33047 is greatly inhibited under light intensities <500 μmol photons m^−2^s^−1^ light.

**Fig 1.**
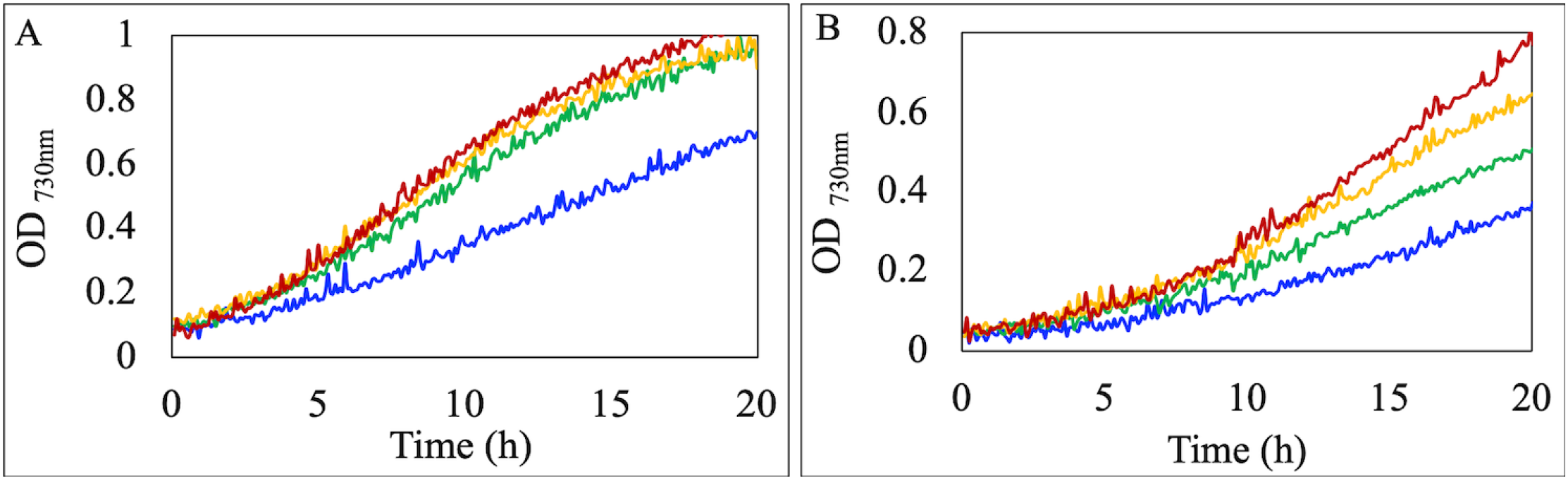
*Anabaena* 33047 thrives under high light. Representative growth curves of WT cells under different light intensities (Blue-500, Green-1000, Yellow-1500, Red-2000 µmol m^−2^ s^−1^). Cells were grown at 42°C in medium with (A) or without (B) added nitrogen sources. Cultures were supplemented with 1% CO_2_.

Light microscopic analysis of filaments grown under nitrogen-fixing conditions revealed higher frequency (14%) of heterocysts in *Anabaena* 33047 as compared to *Anabaena* 7120 (10%) (Fig. S1A) and about 50% higher specific activity of the nitrogenase enzyme when cells were grown under conditions optimal (see methods section) for *Anabaena* 7120 (Fig. S1B).

### Engineering *Anabaena* 33047 for genetic amenability

As anticipated, *the Anabaena* 33047 strain procured from the UTEX culture collection proved to be intractable to all targeted genetic modification attempts. Studies have documented successful transformation of filamentous cyanobacteria by modification of the host restriction modification (RM) system (20). None of the existing conjugal, helper and suicide cargo plasmids (21) yielded any success with conjugation of *Anabaena* 33047. We mined the genome sequence of this strain to ascertain its RM system and subsequently modify it to our advantage. We identified several methylase or methyltransferase genes that appeared to be associated with a type II restriction system (adjacent genes encoded for restriction enzymes). These genes together with their upstream regions were cloned into a helper plasmid (based on helper plasmid pRL623) in various combinations and tested for conjugation efficiency. Various conjugation experiments revealed that a helper plasmid with five genes (see method section for details) in tandem, driven by a lac promoter was most effective in conjugating *Anabaena* 33047 (Fig. S3 A). Using the newly synthesized plasmid, we performed targeted gene deletions and successfully generated a Δ*nblA* mutant of this strain. Surprisingly, unlike other filamentous cyanobacterial strains where single homologous recombination is known to be prevalent in the first few generations, almost all the colonies tested in *Anabaena* 33047 for the gene deletions were double recombinants in the first generation (Fig. S3 B). The complete segregation of the mutant was verified by PCR analysis (Fig. S3 C).

### The *ΔnblA* mutant retains high amounts of PBSs in heterocysts

Fluorescence microscopic analysis revealed that under nitrogen-deficient growth conditions, PBS returned to their normal levels in the vegetative cells of the WT within 24h of onset of nitrogen starvation. However, PBS content of the heterocysts remained significantly lower. In contrast, the Δ*nblA* strain exhibited high amounts of PBS both in the vegetative cells and the heterocysts as the filaments acclimatized to nitrogen deficiency and both cell types appeared brightly fluorescent (Fig. 2 A-D). We quantitated the PBS content in the mutant and WT heterocysts with the help of a fluorescence kinetic microscope (FKM) (Fig. 2 E,F). Steady state fluorescence, depicting the amount of phycobiliproteins in the heterocysts was about eight-fold higher in the Δ*nblA* mutant compared to the WT (Fig. 2G). In contrast, the PBS levels in the vegetative cells of a nitrogen fixing filament was not significantly different between the WT and the mutant strains (data not shown).

**Fig 2.**
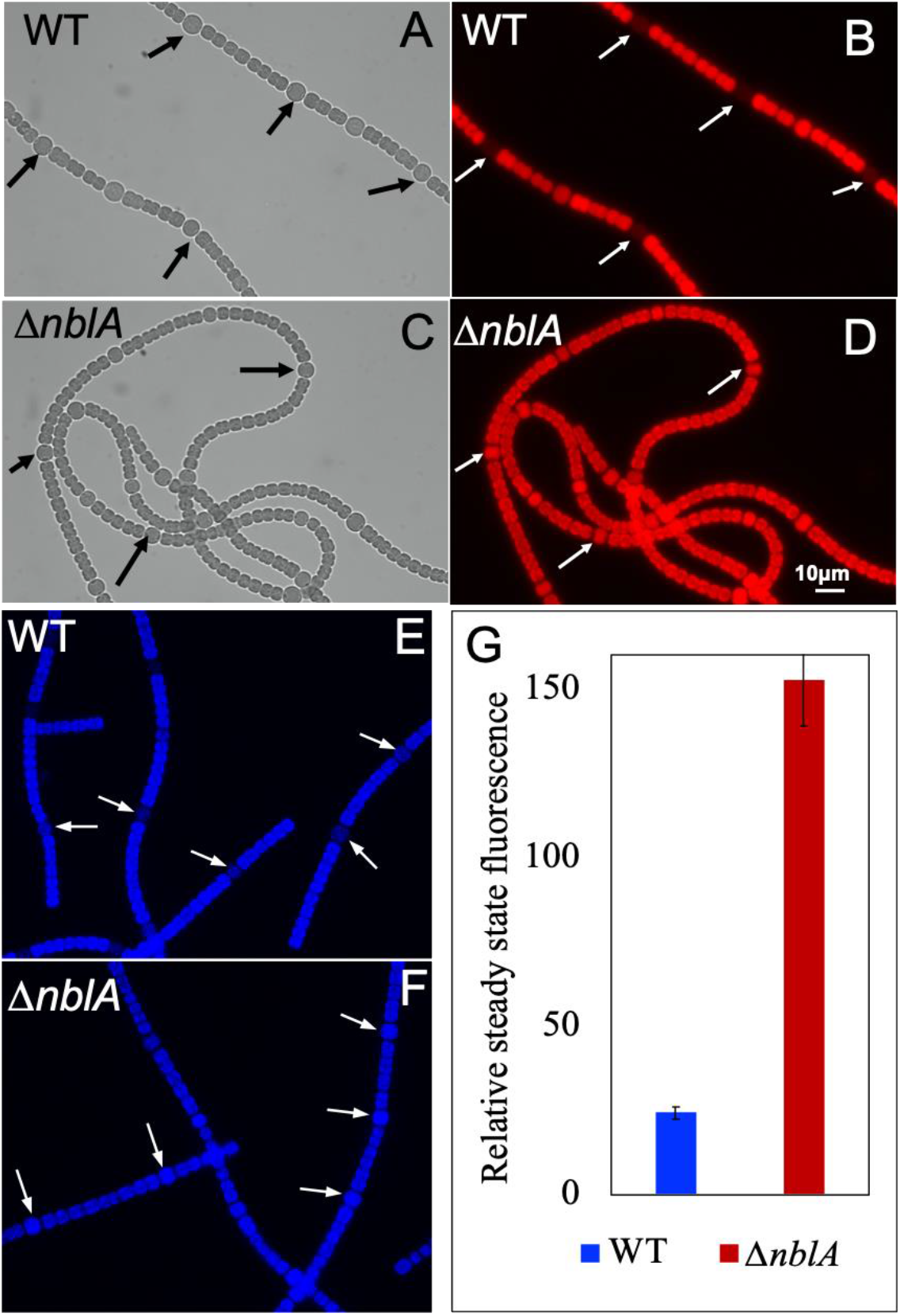
Microscopic analysis showing high amounts of PBS in heterocysts of the Δ*nblA* mutant of *Anabaena* 33047. (A-D) Brightfield and fluorescence microscopic images of WT (A,B) and Δ*nblA* (C,D) mutant of *Anabaena* 33047 grown in media lacking fixed nitrogen sources. High amount of PBS retained in the Δ*nblA* heterocysts compared to the WT (arrows). (E-G) FKM analysis of WT and Δ*nblA* heterocysts. (E,F) Bright signal seen in heterocysts of the Δ*nblA* mutant compared to the WT (arrows). (G) Average of steady state fluorescence obtained from WT and Δ*nblA* mutant after excitation of phycobilisomes in heterocysts.

To evaluate the effect of averting PBS degradation on the physiology of this fast growing, high light tolerant cyanobacterium, we compared the growth rate of the WT and the Δ*nblA* mutant under different light intensities. Differences in growth rate was observed between the mutant and the WT under low light intensities. When grown under 250 μmol photons m^−2^s^−1^, the mutant exhibited ∼50% higher growth rate compared to the WT (Fig. S4). The difference in growth rate narrowed with increasing light intensities and at 500 μmol photons m^−2^s^−1^ the mutant grew ∼ 12% faster than the WT (Fig. S4). The mutant and the WT exhibited comparable growth rates up to 1500 μmol photons m^−2^s^−1^. Under higher light intensities (≥2000 μmol photons m^−2^s^−1^), the mutant initially grew at a slightly reduced rate compared to the WT although final biomass accumulation was similar in both the strains (Fig. S4). The faster growth rate under low light correlated with increased quantum yield and enhanced oxygen evolution rates in the mutant (data not shown) which indicated higher photosynthetic activity. In contrast, photosynthetic activity under high light was slightly reduced.

### The Δ*nblA* mutant exhibits unusually high rates of nitrogenase activity

To assess the effect of PBS retention on heterocyst function in the high light tolerant *Anabaena* 33047, we compared nitrogenase activity in the WT and Δ*nblA* strains. When grown under 250 μmol photons m^−2^s^−1^ light, no significant difference in nitrogenase activity was observed between the mutant and the WT (Fig. 3). However, when grown under 2000 μmol photons m^−2^s^−1^ light, the mutant exhibited 2 to 3-fold higher specific rates of nitrogenase activity compared to the WT (Fig. 3). If grown for more than 36 h under high light in nitrogen deplete medium, both the mutant and the WT filaments clumped into a ball due to excess EPS secretion and both strains exhibited greatly reduced rates of nitrogenase activity. When incubated in the dark, rates of nitrogenase activity were drastically reduced for both the mutant and the WT grown under low or high light intensities (data not shown) indicating that the activity is light dependent.

**Fig 3.**
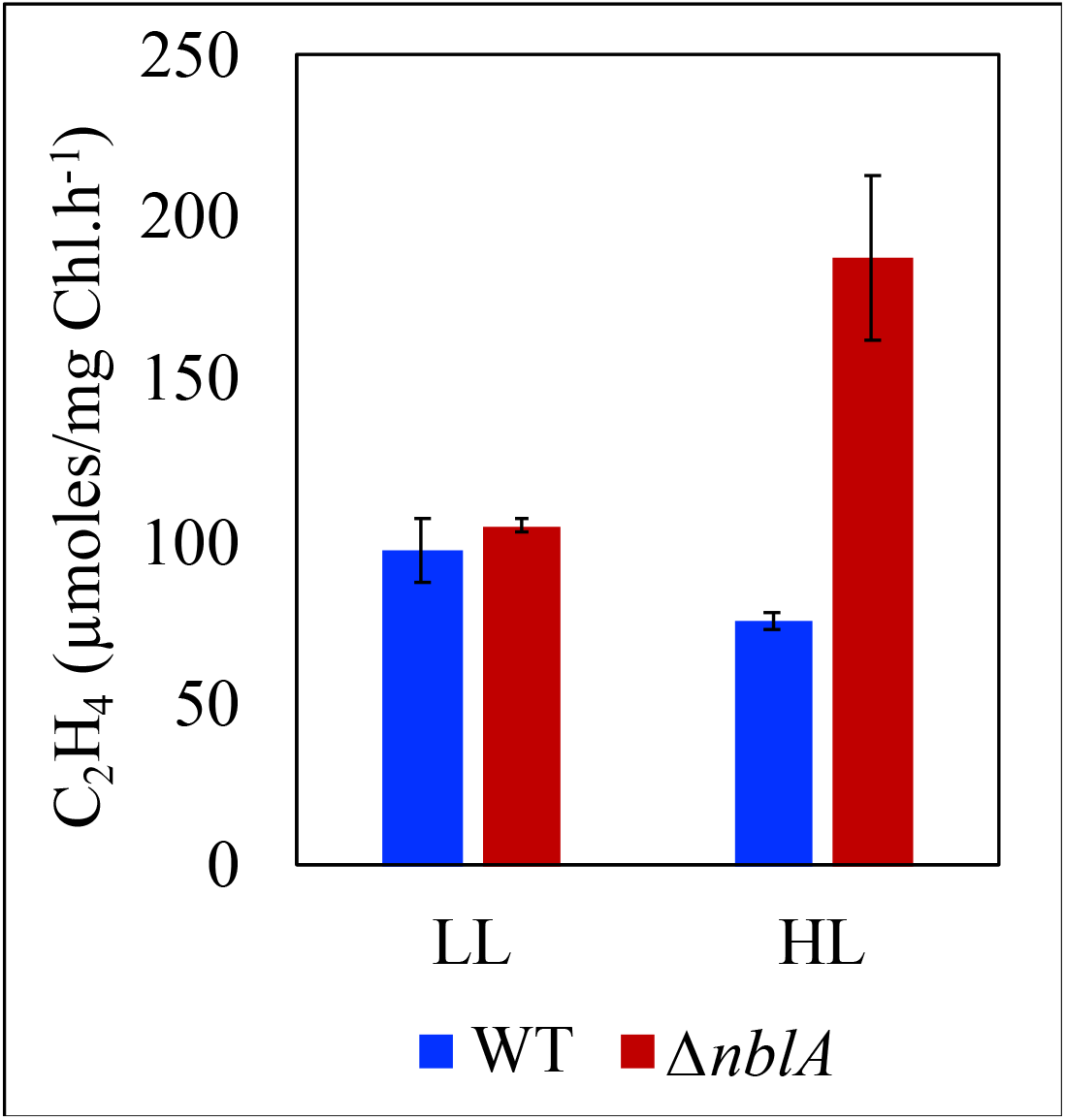
Nitrogenase activity in the WT and Δ*nblA* strains of *Anabaena* 33047 grown in nitrogen deplete medium under low (250 μmol photons m^−2^s^−1^) or high (2000 μmol photons m^−2^s^− 1^) light intensities. Representative data are shown as the average of three biological replicates, and error bars show the standard deviation from the average.

When treated with DBMIB (2,5-dibromo-3-methyl-6-isopropyl benzoquinone), a quinone analogue that inhibits the cytochrome b_6_f complex and thus disables cyclic electron flow(22), nitrogen fixation rates were drastically reduced both in the mutant and the WT. On the other hand, incubation with DCMU (3-(3,4-dichlorophenyl)-1,1-dimethylurea), which blocks PSII and linear electron transport alone, did not have any inhibitory effect on nitrogenase activity in the WT or the mutant (Fig S5).

### P700 oxidation kinetics suggest higher cyclic electron flow in the mutant

To investigate the basis of the enhanced nitrogenase activity in the Δ*nblA* mutant grown under high light and to assess if the mutant exhibited any difference in PSI function, we probed the oxidation/reduction kinetics of PSI reaction centers in both the mutant and the WT using a Joliot type JTS-10 spectrophotometer (23, 24). We exposed dark-adapted WT and Δ*nblA* cells grown under high light to a 5 second pulse of actinic light and measured the oxidation of P700. This was followed by measurement of the re-reduction kinetics in the dark. These measurements were carried out in the absence and presence of the inhibitor DCMU (Fig. S6A, Fig 4), which blocks PSII and linear electron flow in the vegetative cells, thus allowing only cyclic electron flow (CEF) around PSI in the vegetative cells and heterocysts. We also assessed samples treated with DCMU and DBMIB, inhibiting both linear and cyclic electron flows.

**Fig 4.**
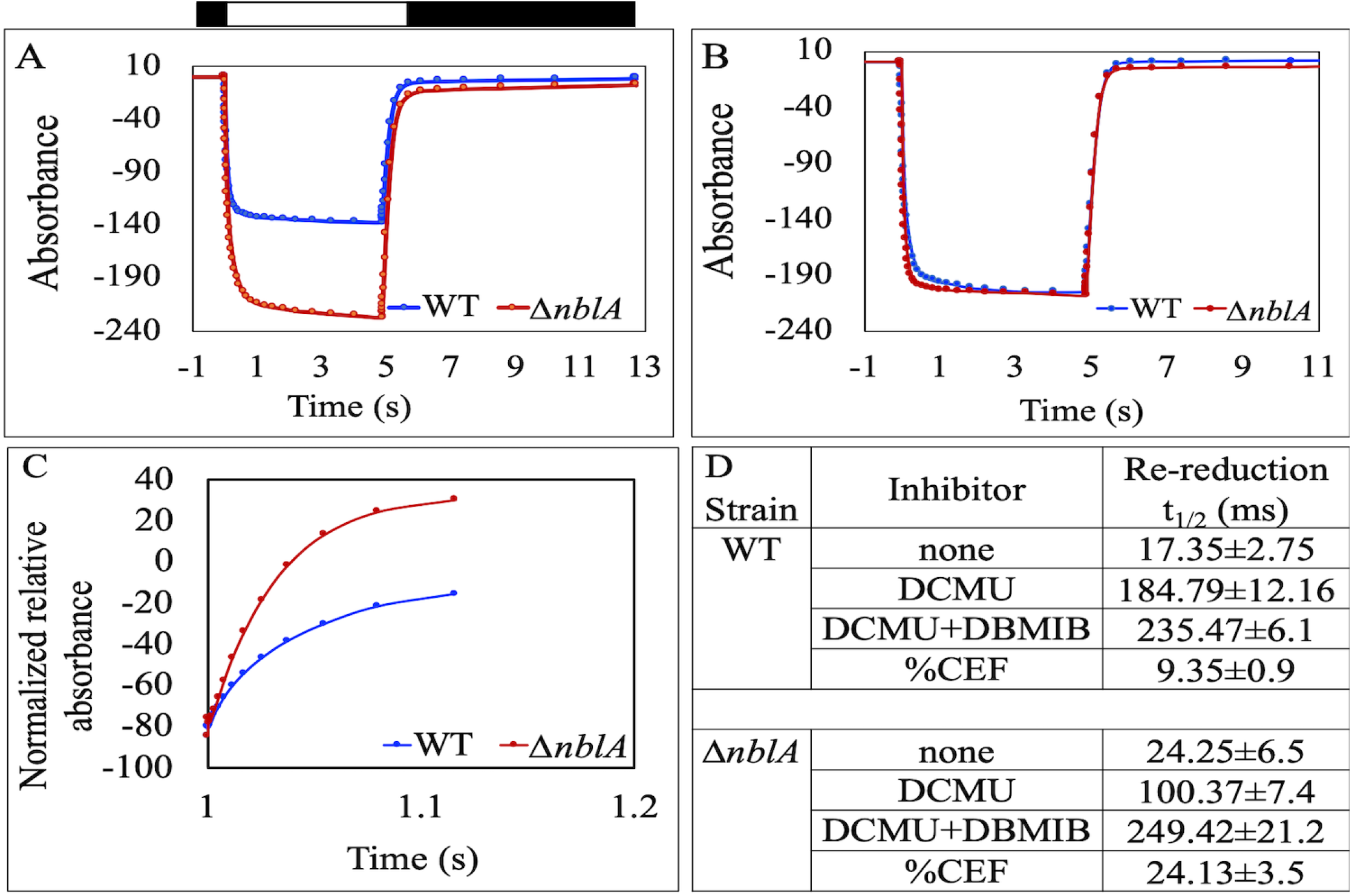
P700 redox kinetics for WT and Δ*nblA* mutant of *Anabaena* 33047 grown under high light (2000 μmol photons m^−2^s^−1^). Dark adapted cells were exposed to a pulse of actinic light for 5 seconds (white bar above panel A). This was followed by dark incubation (black bar above panel A). During this time-course, measuring flashes of 705 nm light probed the redox state of the P700 reaction center of PSI. (A) P700 redox kinetics in DCMU treated (10 μM) WT and Δ*nblA* cells. (B) P700 redox kinetics in WT and Δ*nblA* cells treated with DCMU (10 μM) and DBMIB (1 μM). (C) Details of the re-reduction kinetics of P700^+^ in the dark (normalized to total oxidizable P_700_) for the experiment in panel A. Each trace is an average of 3 independent experiments. (D) The halftimes (milliseconds) for re-reduction of P700 ^+^ in the dark.

When linear electron flow was blocked with DCMU, the P700 oxidation kinetics between the WT and the mutant filaments grown under nitrogen fixing conditions were strikingly different, with greater oxidation of P700 achieved in the Δ*nblA* mutant compared to the WT (Fig. 4 A). In contrast, when treated with DCMU and DBMIB, the oxidation of P700 between the two strains was similar (Fig. 4B). This indicated that the difference observed with DCMU treatment could be a result of enhanced flow of electrons to PSI in the mutant via the cyclic pathway which is obstructed by DBMIB treatment and is not a true reflection of the amount of P700 in the cells. To enable comparison between the mutant and the WT, we normalized the kinetics of P700^+^ re-reduction in the dark (Fig. 4C) to the maximal oxidation observed (24). The Δ*nblA* mutant exhibited faster re-reduction kinetics in the presence of DCMU compared to the WT (Fig. 4C). In contrast, the re-reduction kinetics appeared to be similar in the two strains when treated with DCMU and DBMIB (Fig. S6B). Based on the assumption that CEF is the main contributor to P700^+^ re-reduction, we calculated the percentage contribution of cyclic vs linear electron transport to P700^+^ re-reduction in the WT and mutant filaments (Fig. 4D) (24, 25). When cells grown under high light were treated with DCMU, the mutant showed greater reliance on CEF for P700 oxidation compared to the WT. Under these conditions, the cyclic process accounted for ∼24% electron flow to P700^+^ in the mutant (Fig. 4D). To assess the contribution of vegetative cells to the observed difference in P700 oxidation between the mutant and the WT, we measured P700 oxidation in filaments grown under nitrogen sufficient conditions (filaments without any heterocysts). No significant difference in P700 oxidation was observed between the vegetative cells of the WT and the mutant treated with DCMU (Fig S6C).

## Discussion

*Anabaena* 33047 is unique among heterocystous cyanobacteria in its ability to thrive under very high light intensities. The strain also fixes nitrogen at higher rates (Fig. S1 B) and harbors higher amount of PBS in its heterocysts compared to other heterocystous cyanobacteria (17). This study was initiated to investigate the role of and relationship between high light and PBS content in the heterocysts of *Anabaena* 33047. To this end, we successfully engineered a strategy to modify the genome of this previously recalcitrant strain and generated a *nblA* deletion mutant which retained high amounts of PBS in its heterocysts and enabled us to assess the function of these antenna pigment proteins in nitrogen fixation.

Our current understanding of the effects of averting PBS degradation in cyanobacteria by deleting *nblA* relies largely on studies in non-diazotrophic strains (12, 13, 26–28). The only diazotrophic strain where a Δ*nblA* mutant has been characterized so far is *Anabaena* 7120 and under the conditions tested, the deletion did not have any impact on growth or nitrogen fixation (29). This indicated that degradation of PBS is not an essential adaptive strategy for heterocystous strains to transition from a nitrogen-deplete to a nitrogen-fixing condition.

Various studies have revealed the functional association between PBS and PSI in heterocysts. A study in *Anabaena variabilis* demonstrated a role for PBS in efficient transfer of light energy to PSI and photo-oxidation of P-700, the reaction center of PSI (30). A similar study in *Anabaena* 7120 reported the isolation of a PBS-PSI super complex, the spectral analysis of which revealed efficient energy transfer from PBS to PSI(19). A previously unknown linker component cpcL was shown to be involved in establishing the connection between PBS and PSI. Interestingly, this linker is found in many heterocystous cyanobacteria, including *Anabaena* 33047, but not in unicellular cyanobacteria that fix nitrogen at night. The study found that the levels of this linker were four-fold higher in heterocysts compared to vegetative cells and proposed a role for the PBS-cpcL-PSI complex in harvesting light energy and facilitating PSI driven nitrogen fixation. These findings were also supported by a more recent study which showed transfer of energy from cpcL-PBS to PSI in a mutant of *Anabaena* 7120 (4). In addition, a role for these phycobiliproteins in supporting light-driven nitrogenase activity in heterocysts was also demonstrated (30).

In heterocystous cyanobacteria cyclic electron flow around PSI plays a crucial role in driving nitrogenase activity (31–33). Cyclic electron flow relies on the transfer of reducing equivalents from PSI to the plastoquinone pool via the NDH-1 complex. The reduced plastoquinone pool is re-oxidized by the cytochrome-b6/f (Cyt-b6/f) complex which transfers electrons back to PSI via plastocyanin, thereby completing the cycle. This cyclic flow of electrons is coupled to the generation of a proton gradient across the thylakoid membrane which drives ATP synthesis (Fig. 5). CEF around PSI is light dependent and it has been demonstrated that CEF can vary with light intensity, with increased irradiance leading to higher rates of CEF (34, 35).

**Fig 5.**
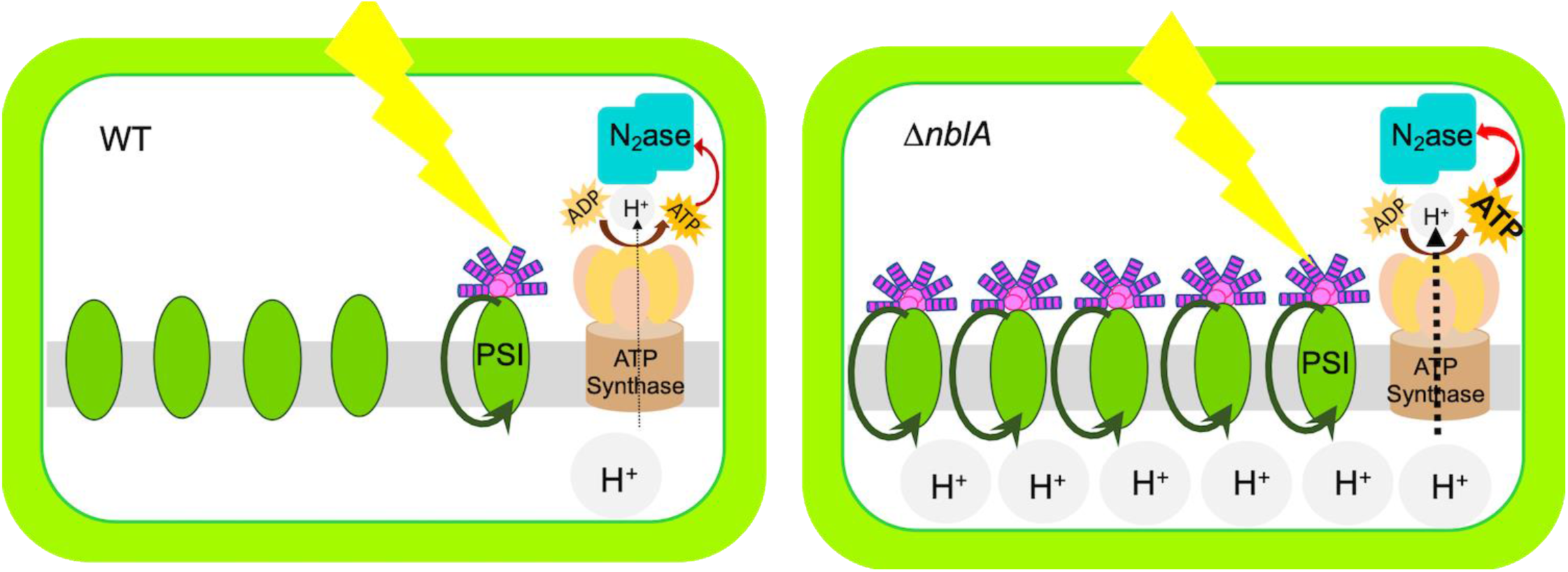
Schematics depicting the differences in the heterocysts of the WT and Δ*nblA* strains that contribute to enhanced nitrogenase activity. Higher abundance of phycobilisomes in the mutant heterocyst and their association with PSI centers leads to higher ATP generation mediated by cyclic electron flow. The ATP feeds into the nitrogenase enzyme complex leading to higher rates of nitrogen fixation in the Δ*nblA* mutant.

The Δ*nblA* mutant exhibited higher PBS content in its heterocysts and higher nitrogenase activity. Spectroscopic studies demonstrated higher CEF in the mutant compared to the WT (Figs. 2, 3, 4). Our FKM analysis did not reveal any significant difference in pigment content between the vegetative cells of the mutant and the WT (after PBS is resynthesized in the WT) and no significant difference was observed in P700 oxidation between the vegetative cells of the mutant and the WT grown under nitrogen sufficient conditions. These observations suggest that the enhanced CEF detected in the mutant is likely a reflection of the altered photochemistry in its heterocysts brought about by higher levels of PBS which in turn contributes to enhanced nitrogenase activity. In addition, averting PBS degradation can eliminate the need for the expensive re-synthesis of these large antenna complexes, a phenomenon that likely takes place in the wild type *Anabaena* 33047 heterocysts that harbor higher amounts of PBS compared to other cyanobacteria. Averting antenna degradation in the Δ*nblA* mutant possibly allows utilization of the cellular resources for the energy-intensive nitrogen fixation process instead. Retention of PBS in the vegetative cells of the mutant during acclimation to nitrogen limitation did not seem to have any significant adverse effect on growth or metabolism. A small inhibitory effect on growth was observed for a few hours after inoculation into nitrogen deplete medium which could be a result of a slight reduction in photosynthesis observed under these conditions, but the final biomass accumulation was similar to the WT. Interestingly, the higher nitrogenase activity in the mutant did not contribute to faster growth under high light, suggesting that other cellular resources could be limiting or that the high rates of N_2_-fixation achieved by the WT are optimal to support fast growth of this strain. The excess N_2_ in the Δ*nblA* strain is probably channelized into storage reserves. When grown for an extended period of time (>30h) excess EPS secretion led to clumping of the filaments into a tight ball thereby limiting light availability to the cells resulting in greatly reduced nitrogenase activity indicating that access to high light is crucial for augmenting nitrogenase function. In contrast, when grown under low light, the presence of PBS turned out to be advantageous for efficient light harvesting in the mutant and this was reflected in faster growth compared to the WT(Fig. S4). However, the low light growth conditions could not elicit an increase in nitrogenase activity. This again suggests that the high rates of nitrogenase activity in the WT *Anabaena* 33047 is optimal to support growth under all conditions but light can be a limiting factor4. The mutant thus holds the potential to channelize the excess pool of nitrogen fixed under high light towards nitrogen-rich products of interest.

Our study demonstrates a link between PBS content and nitrogenase activity in heterocysts. Increased light absorption by PBS leads to enhanced cyclic phosphorylation which is the driving force for nitrogen fixation. Thus, in a high light tolerant strain like *Anabaena* 33047 modifying the heterocyst antenna by disabling degradation of PBS during nitrogen acclimation appears to be an effective strategy for enhancing nitrogen-fixation rates.

## Materials and Methods

### Cyanobacterial strains and growth conditions

The *Anabaena* sp. ATCC 33047 strain was procured from the UTEX Culture Collection of Algae at the University of Texas at Austin (www.utex.org). The strain was isolated from the Texas Gulf coast more than five decades ago. It was then designated as *Anabaena* CA (36,37) For conjugation experiments, cells were grown in BG11 medium with added nitrate, in shake flasks (∼150 rpm), under 150μmol photons m^−2^s^−1^ of white light in ambient air at 38°C. For physiological studies, cells were grown in ASP2 liquid medium with or without added nitrate at desired light intensities. The *Anabaena* sp. ATCC 7120 strain was also acquired from the UTEX Culture Collection and cells were grown in BG11 medium at 50μmol photons m^−2^s^−1^ of white light in ambient air at 28°C with or without added nitrate.

### Genetic modification and construction of strains

The *ΔnblA* strain was generated by replacing the *nblA* gene with a kanamycin resistance cassette using homologous recombination. The deletion construct was conjugated into *Anabaena* 33047 using a modified helper plasmid (pSL3348), containing five methylase or methyltransferase genes from the genome of *Anabaena* 33047 (Supplementary table 1). Details of vector construction and conjugation protocol are provided as supplementary information.

### Growth Analysis

For growth measurements, *Anabaena* 33047 cultures were maintained in ASP2 medium, with shaking at ∼150rpm, under 250μmol photons m^−2^s^−1^ of white light in ambient air at 38°C. *Anabaena* 7120 cultures were maintained in BG11 medium, with shaking at ∼150rpm, under 50μmol photons m^−2^s^−1^ of white light in ambient air at 28°C. Culture aliquots were then diluted to an OD730 of 0.05 in a Multi-Cultivator MC 1000-OD device (Photon Systems Instruments, Drasov, Czech Republic), and growth under continuous light-emitting diode (LED) light of different intensities in ambient air supplemented with 1% CO_2_ were measured at 730nm.

### Fluorescence Microscopy

Cells from 24-48h liquid culture were imaged using a Nikon Eclipse 80i microscope equipped with a Photometrics Cool Snap ES CCD camera (Roper Scientific). Illumination was provided by a metal halide light source (X-Cite). PBS fluorescence was detected using a 560/40nm excitation filter, a 595nm dichroic beam splitter, and a 630/60nm emission filter.

### Measuring PBS in heterocysts

Steady state fluorescence kinetics of WT and Δ*nblA* heterocysts of *Anabaena* 33047 were measured by the Fluorescence Kinetic Microscope (Photon Systems Instruments, Drasov, Czech Republic, www.psi.cz). PBS were excited with green LED light (LZ1-00G100, 530 nm peak). A steady state PBS fluorescence was separated from exciting light by a dichroic mirror with 562nm edge wavelength (Semrock 562nm edge BrightLine^®^) and collected using an emission filter with transmission band 663.5 – 666.5 nm (Semrock 660/13nm BrightLine^®^). Signal were measured from individual heterocysts and vegetative cells using the 40x objective (Zeiss Plan-Apochromat 40x/1.4 Oil). The average signal from a minimum of 10 cells was used to obtain the representative PBS signal from the mutant and the WT. The FluorCam7 software developed by Photon Systems Instruments was used to operate the FKM and analyze the data.

### Nitrogen fixation assay

Nitrogenase activity was measured using the acetylene reduction assay, as described in reference (38) and expressed in terms of the ethylene produced per mg of chlorophyll. Cultures were grown in ASP2 medium lacking fixed nitrogen in multicultivator tubes under 250 and 2000 μmol photons m^−2^s^−1^, at 42°C in air supplemented with 3% CO_2_. Cells were transferred to airtight 100ml glass vials and incubated in a 5% acetylene atmosphere under light at 450μmol photons m^−2^s^−1^ 38°C for 3h. Gas samples were withdrawn from the vials, and ethylene production was measured using an Agilent 6890N gas chromatograph equipped with a Poropak N column and a flame ionization detector, with argon as the carrier gas (38).

### PSI measurements

P700 concentration in whole cells was determined using a JTS-10 pump probe spectrometer (BioLogic, France). Cultures grown to mid-exponential phase were normalized to 5 μg/mL chlorophyll and incubated at 38°C under light prior to measurements. For analysis of P700 kinetics, samples were treated with the required inhibitors (10µM DCMU, 20µM DBMIB) and dark adapted for 3 min before subjecting them to an actinic light pulse. The redox state of P700 was monitored by absorption at 705 nm during the 5 min saturating pulse and for 10 seconds of recovery afterwards.

## ACKNOWLEDGEMENTS

This study was supported by funding from the U.S. Department of Energy, Office of Science, Office of Biological and Environmental Research, Genomic Science Program under Award Number DE-SC0019386 and Gordon and Betty Moore Foundation (GBMF5760) to HBP. ZY’s visit to Washington University was also supported by the National Natural Science Foundation of China (31400215).

## Supplementary information - legends

### Figures

Figure S1 Characterization of *Anabaena* 33047. (A) Comparison of bright field and fluorescence images of filaments of *Anabaena* 33047 and *Anabaena* 7120 grown under nitrogen fixing conditions. Higher frequency of heterocysts is observed in *Anabaena* 33047 (arrows). (B) Comparison of nitrogenase activity in *Anabaena* 33047 and *Anabaena* 7120. Representative data are shown as the averages of three biological replicates, and error bars show the SDs from the averages.

Figure S2 Representative growth curves showing comparison of WT *Anabaena* 7120 grown under 400 µmol m^−2^ s^−1^ light intensities at 28°C in medium with (blue) and without (green) sources of fixed nitrogen and supplemented with 1% CO_2_.

Figure S3 Engineering genetic amenability in *Anabaena* 33047. (A) Schematics of the strategy used for designing the helper plasmid that was used in tri-parental conjugation of *Anabaena* 33047. (B) Segregation analysis of the Δ*nblA* mutant. (C) PCR analysis verifying segregation of the Δ*nblA* mutant.

Figure S4 Representative curves showing growth comparison of WT (solid lines) and Δ*nblA* (dotted lines) strains of *Anabaena* 33047 under low and high light intensities at 42°C in medium without sources of fixed nitrogen and supplemented with 1% CO_2_. Red - 2000 µmol m^−2^ s^−1^, green - 500 µmol m^−2^ s^−1^, grey - 250 µmol m^−2^ s^−1^.

Figure S5 Nitrogenase activity in the WT and Δ*nblA* strains of *Anabaena* 33047 grown in nitrogen deplete medium and assayed in the presence of the inhibitors DCMU (10 μM) and DBMIB (1 μM). Representative data are shown as the average of three biological replicates, and error bars show the standard deviation from the average.

Figure S6 P700 redox kinetics for WT and Δ*nblA* mutant of *Anabaena* 33047 grown under high light (2000 μmol photons m^−2^s^−1^). (A) P700 kinetics in the WT and Δ*nblA* mutant in the absence of any inhibitor. (B) The details of the re-reduction kinetics of P700^+^ in the dark (normalized to total oxidizable P_700_) for the experiments in figure 4B. (C) P700 kinetics in the WT and Δ*nblA* mutant grown under nitrogen sufficient conditions and assayed in the presence of DCMU (10µM). Each trace is an average of 3 independent experiments.

### Table

**Table** S1 Details of the Methylase or methyl transferase genes and their upstream regions that were cloned into the newly constructed helper plasmid pSL3348 that was used to successfully conjugate *Anabaena* 33047 and generate the Δ*nblA* mutant.

### Text

**Materials and Methods** Genetic modification and construction of mutant

